# Isolation of Plastic Digesting Microbes from the Gastrointestinal Tract of *Tenebrio Molitor*

**DOI:** 10.1101/2024.10.16.618709

**Authors:** Hannah McKinnon Reish, Rebecca F. Witty, Adam H. Quade, Jason W. Dallas, Lucas J. Kirschman

## Abstract

Plastics, particularly polystyrene (PS) and polyethylene (PE), are pervasive in ecosystems and degrade slowly, leading to the formation of microplastics and nanoplastics, which pose environmental and health risks. The gut microbiota of *Tenebrio molitor* larvae (mealworms) can degrade plastics, making them a potential model for studying microbial plastic digestion. This study investigated how rearing *T. molitor* on PS-or PE-supplemented diets over three generations affected gut microbiota composition and plastic degradation. Larvae were reared on wheat bran (control), PS, or PE diets, and microbial communities from their guts were cultured to assess plastic metabolism. Results showed that the PE-supplemented diet enhanced bacterial plastic degradation, while the PS-supplemented diet did not lead to similar improvements. Microbes from PE-fed larvae exhibited increased plastic metabolic activity compared to controls, and several bacterial strains with plastic-degrading capabilities were identified. However, microbial digestion of PS was less efficient, and potential dysbiosis due to the presence of *Dickeya* phage in PS-fed larvae may have influenced the results. Further research is needed to clarify the mechanisms driving microbial plastic digestion and to determine the applicability of these findings to environmental plastic remediation, particularly for micro– and nanoplastics.

## INTRODUCTION

The presence of plastics is ubiquitous in aquatic and terrestrial ecosystems and is considered an indicator of the Anthropocene [1]. Worldwide plastic production was about 400 million metric tons in 2022 and will increase exponentially [2] with the United States, alone, producing up to 40 billion metric tons by 2050 [1]. Recovery and recycling of these plastic products trails behind its production almost tenfold [3]. Global estimates in 2017 indicated that about 8300 million tons of virgin plastics had been produced, with less than 10% of that being recycled and over 75% accumulating in landfills and natural environments [4]. Current plastic production is exponentially outpacing recycling or sequestration capacity, and only a small subset of plastics can be recycled into new consumer products at scale [5–8]. Furthermore, physical and chemical degradation of plastic waste can give rise to microplastics (particles < 5mm) and nanoplastics (particles < 1,000 nm) [9]. These particles are ubiquitous, particularly in marine environments. Beaches typically have 500 – 27,000 microplastic particles per square meter, ocean water can have up to 15,000 particles per cubic meter, and ocean sediment can contain 2,000 particles per kilogram [10]. The amount of microplastics in terrestrial systems varies widely, as non-agricultural soils may be nearly microplastic free but agricultural soils can contain up to 42,000 particles per kilogram [11]. Both microplastics and nanoplastics can cause inflammation, oxidative stress, and toxicity [9,12–14]. For example, microplastics and nanoplastics have been found inside human carotid artery plaques where they increased the risk of cardiovascular events and death [15]. The scale of plastic production and the threats posed by micro– and nanoplastics have led to a ‘plastic panic’ resulting in bioprospecting for novel biodegradation routes as an option for remediation [8].

Polystyrene (PS) and polyethylene (PE) make up most single-use plastic consumer products and packaging [16,17]. Polystyrene is a long chain molecule comprised of smaller styrene monomers of styrene (C_8_H_8_). Polystyrene is a parent polymer of styrene-based plastics since it was developed in the 1950’s [4,18]. It has a wide range of both commercial and industrial uses, from food safe storage, insulation and furniture, and may contain additives or co-monomers depending on purpose of the end products [18]. Polyethylene is a polymer comprised of several small molecules of ethylene (CH_2_). Polyethylene may also contain additives dependent on its commercial or industrial purpose. These additives can include antioxidants, pigments, rubbers, or flame retardants [19].

Both PS and PE are long strand chains that are resistant to many forms of degradation [20,21]. Due to their high molecular weight, stable structure, and hydrophobicity, both plastics have long environmental half-lives and reprocessing is only possible about 20 times before their mechanical or thermal properties change [18,22,23]. Researchers have shown levels of biodegradation of high molecular weight PE in soil, although there is no consensus as to how long complete biodegradation actually takes with the estimates ranging between 10 and 32 years [24,25]. Currently, only intense thermal treatments like combustion or pyrolysis are seen as permanent methods to eliminate plastics, but these methods are incredibly energy intensive and expensive, making them and unrealistic solution to the growing issue [4].

Scientists have found evidence of natural plastic degradation in many surprising sources ranging from marine polychaetes to soil samples [26,27]. The majority of current research focused on the characterization of the microbes in these systems. Some studies have explored the degradation by microbes, but results have not been promising, with outcomes of 0.8% of gravimetric weight loss over an 8 week incubation, or a maximum efficiency of 10% when degrading a PS composite [21,28,29]. Researchers are working to engineer solutions by developing strains of *Escherichia coli* or *Pseudomonas* sp. that can upcycle or break down different plastics via a *de novo* pathway or enzyme transformation [30,31]. A third avenue, entomoremediation, has risen in popularity, with several species of insects being observed to chew through several plastic types while in their larval form [32]. Most notably darkling beetle larvae (*Tenebrio molitor* and *T. obscurus*) have been found to host PS and PE digesting microbes, *Citrobacter sp.* and *Enterobacter sp.* in their gut microbiome [33]. This observation has been confirmed by researchers and citizen scientists across 20+ countries, with fecal analysis confirming cleavage of the long-chain PS hydrocarbons [8,20,27]. However, insects, *per se*, are not a viable solution to the plastic problem as they require too much carbon input, and current methods have not streamlined the processes to be cost effective [33,34]. Polystyrene and PE alone provide only hydrocarbons, they do not supply *T. molitor* with protein or micronutrients [20]. Plastic supplementation alone results in larvae cannibalizing each other and consuming molts for nutrients [20,35]. However, selection for and isolation of gut microbes responsible for plastic biodegradation may be a promising alternative that avoids the downsides of entomoremediation.

With respect to entomoremediation, research on the gut microbiome of *T. molitor* has mainly focused on characterization of the bacterial community [36,37]. One such study found that in a single generation, the microbial community underwent a significant shift when the diet was changed from wheat bran to plastic (PS or PE) and then back to wheat bran, over consecutive two week periods [38]. To our knowledge, no experiment has exceeded two generations of sampling which hinders our understanding of how long-term plastic exposure influences the gut microbiome. Here, we aimed to observe how plastic inclusion in the diet of *T. molitor* over three generations would alter the larvae’s microbial ability to biodegrade plastics. Based on a preliminary finding where the F_1_ generation larvae exhibited improved ability to digest plastic, compared to their P generation [39], we examined whither three generations would improve plastic digestion by isolated gut bacteria.

## METHODS

### Insect Husbandry

We haphazardly sorted about 9000 *T. molitor* larvae (Timberline Live Pet Foods) into three separate treatments: control, PS supplemented, and PE supplemented. Each treatment contained about 3,000 larvae that represented the P generation. The larvae were housed in stacked drawers (32.1 cm × 25.4 cm × 17.8 cm; Sterilite Corp.) and each drawer within a treatment separated life stage (larvae, pupae, adults). The adult drawer for each treatment had a wire mesh floor, which enabled recently hatched larvae to fall through into the drawer below, thus separating generations. We manually removed pupae from the larval drawer and placed them in their own dedicated drawer. When the beetles pupated, they were placed in a new adult drawer. Each drawer was covered in black duct tape to minimize light. We maintained ambient temperature between 24 – 26 °C, ∼50% humidity, on a 12-hour light cycle.

Substrates were treatment specific. The control population substrate was wheat bran, and the two experimental populations were reared on a 5:1 wheat bran to plastic ratio (PS and PE). We shredded PE foam (Foam Factory Inc., 1.27 cm thickness, 0.82 kg density) and low-density PS foam (Foam Factory Inc., 2.47 cm thickness, 0.45 kg density) with a food processor (General Electric, model: G8P1AASSPSS). Because of the mass differences in the plastics and bran, the treatments did not receive the same mass of food, rather we kept the food at a constant level in the drawers ∼5 cm below the top. We provided slices of potato daily as a water source.

### Microbial Culture

At the end of the F_3_ generation, we randomly selected 100 larvae from each treatment and fed them a plastic only diet for one week. After a week, the gastrointestinal tracts of 50 larvae per treatment were extracted with sterilized, dedicated equipment. Larvae were euthanized and surface sterilized in 90% EtOH, and their GI tracts were homogenized in a 0.75% sterile saline solution. The resulting product was then used to establish gut cultures in triplicate.

Cultures were incubated in 30 mL Bushnell-Haas broth (MG Scientific; #257820). Bushnell-Haas broth is carbon-free, making the addition of either PS or PE film the only carbon source. For the cultures incubated with PE, we added 3 g of commercially obtained plastic film (Topco Associates LLC, Simply Done Plastic Wrap, SN: 011225005022). For the cultures that we incubated with PS, we added 3 g of PS film created based on the methods of Yang et al. [40]. Briefly, we dissolved PS in a xylene solvent (0.03 g mL^-1^) and spread the resulting solution over glass. We removed the film after 24 hours and then dried it in a fume hood for 72 hours at room temperature. We then rinsed the films with methanol solvent and deionized water to remove residual xylene from the films.

We incubated cultures from control larvae on PS and PE. Cultures from larvae raised on PE or PS diets were incubated with the corresponding plastic type. We also included process controls, which contained no microbial cultures were incubated with each plastic type. Culture tubes were stored in a shaker (150 rpm) (New Brunswick Scientific, C24) at 37 °C for 60 days. Every 7-10 days 10 ml of media was removed and disposed of, and 10 mL of fresh media was added. After 60 days the resulting cultures were spread on nonselective tryptic soy agar (TSA) plates and incubated overnight at 37 °C.

### Microbial Isolation, Metabolism, and Sequencing

Microbial isolation and measurements of microbial plastic metabolism followed the methods of Brandon et al. [41] and Yang et al. [27]. First, we isolated strains via morphology on TSA and then resuspended isolates in tryptic soy broth and cultured them for 48 hours at 37 °C. We then centrifuged these cultures at 5000 rpm for five minutes, pipetted off the liquid, added 2 mL of 0.75% sterile saline solution, and vortexed for 30 s to resuspend the microbes. We repeated this process three times. Following the final resuspension, we diluted all the samples to 0.9 OD_600_.

The microbial dilutions were added, in triplicate, to a Biolog MT2 96-well Microplates (Hayward). MT2 microplates contain a carbon-free nutrient medium and tetrazolium violet, a redox dye. This is an established indicator of bacterial carbon oxidation and has been used previously to assess plastic oxidation by bacteria from *T. molitor guts* [41–43]. The three replicate wells for each bacteria contained ∼3 mg microplastics, PS (PSPM-1.07, 85-105 μm, Cospheric) or PE (CPMS-0.96, 125-150 μm, Cospheric). We included positive controls containing ∼3 mg glucose (G-8270, Sigma, St. Lois), negative controls that contained only saline and bacteria, and template controls that contained only microplastics and saline. We measured the absorption every hour over 24 hours at 595 nm on a Multiskan FC Microplate Spectrometer (Fisher Scientific). The samples were automatically stirred between measurements. From the resulting measurements, we calculated the slope of the linear phase of the reaction. And determined the measurement at hour twelve was peak metabolism. We normalized all the values by subtracting the plastic refraction from the template controls and then corrected each isolate by subtracting the negative control from each replicate. This left bacterial metabolism as a normalized absorbance (595 nm) as our final value [41].

We sent TSA cultures of the 20 strains with the highest normalized absorbance (595 nm) for identification via 16S rRNA gene sequencing at Azenta Life Sciences Inc. Submitted colony samples underwent a crude NaOH lysis to be directly used in PCR. PCR was performed using PCR primers that amplify regions V1 – V9 of the 16S rRNA gene, resulting in an amplicon approximately 1.4 kb in size. Colonies that were run on the eukaryote ITS assay were targeted with ITS1 and ITS4 primers. Amplified samples were spot checked using gel electrophoresis to check for robust amplification along with a negative control to check for contamination. Following amplification, enzymatic cleanup was performed using Exonuclease I – Shrimp Alkaline Phosphatase (ExoSAP). Dye-terminator sequencing Applied Biosystems BigDye version 3.1. Sequencing-specific primers were used to generate bidirectional reads. The reactions were then run on Applied Biosystem’s 3730xl DNA Analyzer. If applicable, bacterial identification was performed by using an in-house bioinformatics algorithm. The script compares de novo assemblies of each sample to NCBI’s BLASTn database. The database used is RefSeq 16S. An E-value threshold of 1e-250 is used for reporting. The best/most specific genus or species is provided in the report. BLAST identification is not performed by Azenta for eukaryote ITS assays.

### Gut Microbiota 16S Characterization

During each generation, we randomly selected 10 larvae from each treatment group. We euthanized and surface sterilized the larvae in 95% EtOH before removal of their GI tracts with dedicated, sterilized equipment. The guts were then flash frozen with liquid nitrogen and stored at –80°C. Samples were sent to CD Genomics for 16S rRNA gene sequencing. 16S rRNA gene sequencing was performed on the Illumina MiSeq platform using a paired-end 300 bp approach. Barcodes and primer sequences were removed from the raw reads, which were then merged using FLASH (v1.2.11) to produce raw tags. Quality filtering was applied using Fastp to remove low-quality reads, yielding high-quality clean tags. The sequences were further processed using QIIME2, where the DADA2 algorithm was employed to generate Amplicon Sequence Variants (ASVs).

### Statistical Analyses

Statistical analyses of our results are constrained due to pseudoreplication arising from our experimental design. Each of the three larval treatments had larvae housed together within their respective treatment, and larval guts were pooled before incubation with the plastics and subsequent microbial isolation. This lack of independent experimental units means that treating samples as independent replicates could lead to misleading conclusions and increased Type I error rates [44–47]. All analysis were performed in R (version 4.3.1; R Project for Statistical Computing) with R Studio (version 2023.06.1; PositTM).

To assess patterns in how gut microbiota changed between treatments while acknowledging the potential confounding effect of co-housing, we employed multivariate techniques. With respect to the recommendations of the American Statistical Association’s recommendations, we did not set an alpha level for significance [48]. Prior to analysis, we rarefied the data using the ‘rrarefy’ function in the vegan package [49]. We performed non-metric multidimensional scaling (NMDS) using the Bray-Curtis dissimilarity matrix to visualize patterns in multivariate space. The betadisper function from the vegan package did not show multivariate homogeneity of groups dispersions (variances). Five NMDS ordinations, beginning with one dimension, were created using 10,000 permutations and compared for appropriate stress values (k < 0.20) and dimensionality suing the metaMDS function of the vegan package. We then conducted a permutational MANOVA (PerMANOVA) using the Bray-Curtis dissimilarity matrix and 10,000 permutations to test for differences between treatments. Permutational pairwise comparisons were performed using the RVAideMemoire package [50]. Given the pseudoreplication in our design, we acknowledge that any observed differences could be attributed to either treatment effects or the co-housing of larvae within treatments. We attempted to account for this by using a blocking factor in the PerMANOVA; however, because there was only one housing unit per treatment due to the co-housing, the model was unable to partition the variance appropriately and returned a p-value of 1.0. Therefore, we ran the PerMANOVA without the blocking factor. We also examined the abundance of bacteria by class and computed Shannon-Weiner indices for each larval community from OTUs. The results of the microbial respiration assay were analyzed as the mean (± SD) for the triplicate assays for each microbial isolate. We removed any microbe with a replicate < 0 from the analysis.

## RESULTS

### Gut Community Structure

We detected 4,186 ASVs from 25 phyla, 45 classes (Fig. 1), 170 families, and 275 genera. Several unique, unassigned phyla were detected, as well as a *Dickeya* phage that contains a 16S gene. As a bacteriophage, *Dickeya* phage has a predatory role as a virus specialized to infect *Dickeya sp.*[51]. The *Dickeya* phage was highly abundant in both experimental treatments, with 30% relative abundance in the PS samples and 12% in the PE samples, but it was in less about 1% of both the Parental and Control groups (Fig. 1a). After removing *Dickeya* phage, and recalculating percentages, Firmicutes was the most abundant phylum in each treatment (parental: 83%, control: 88%, PE: 97%, PS: 89%) with Bacilli being the most prevalent class (parental: 84%, control: 76%, PE: 93%, PS: 64%).

**Figure 1.**
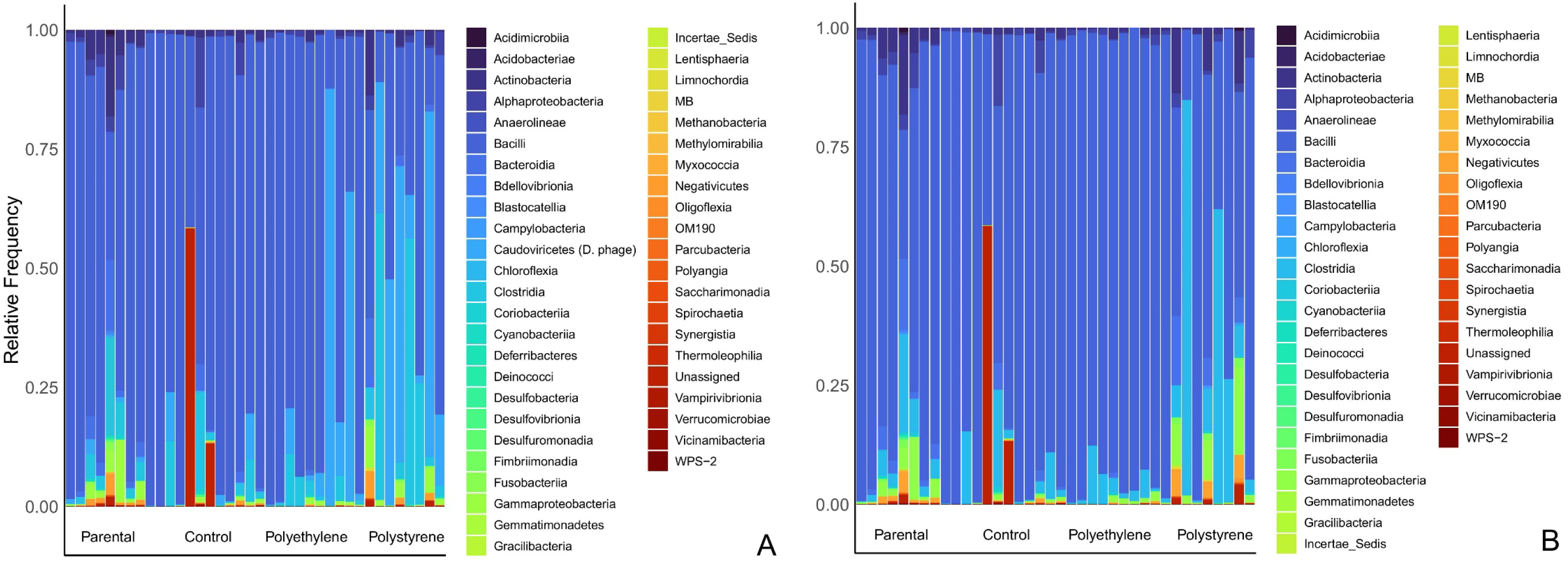
Relative abundance of bacteria classes isolated from Tenebrio molitor larvae for the parental generation and three treatments at the F_3_ generation. The Caudoviricetes class, which was represented by the Dickeya phage, is present in A and removed in B.

Ordination and PerMANOVA analyses (Table S1 & S2) suggest differences in the gut microbiota community both with *Dickeya* phage (Fig. 2A; k= 3, stress = 0.12, R^2^ = 0.22, df = 3, F = 3.3. p < 0.0001) and without *Dickeya* phage (Fig. 2B; k= 3, stress = 0.11, R^2^ = 0.2, df = 3, F = 3, p = 0.0007). The observed patterns and differences remained consistent after the removal of the bacteriophage. We observed pairwise differences between parental and control groups (p = 0.02 with bacteriophage, p = 0.02 without) and between parental and PS (p = 0.002 with bacteriophage, p = 0.002 without), but not between parental and PE (p = 0.5 with bacteriophage, p = 0.6 without). We also found differences between Control and PE (p = 0.02 with bacteriophage, p = 0.03 without), Control and PS (p = 0.005 with bacteriophage, p = 0.006 without), and between PE and PS (p = 0.02 with bacteriophage, p = 0.02 without).

**Figure 2.**
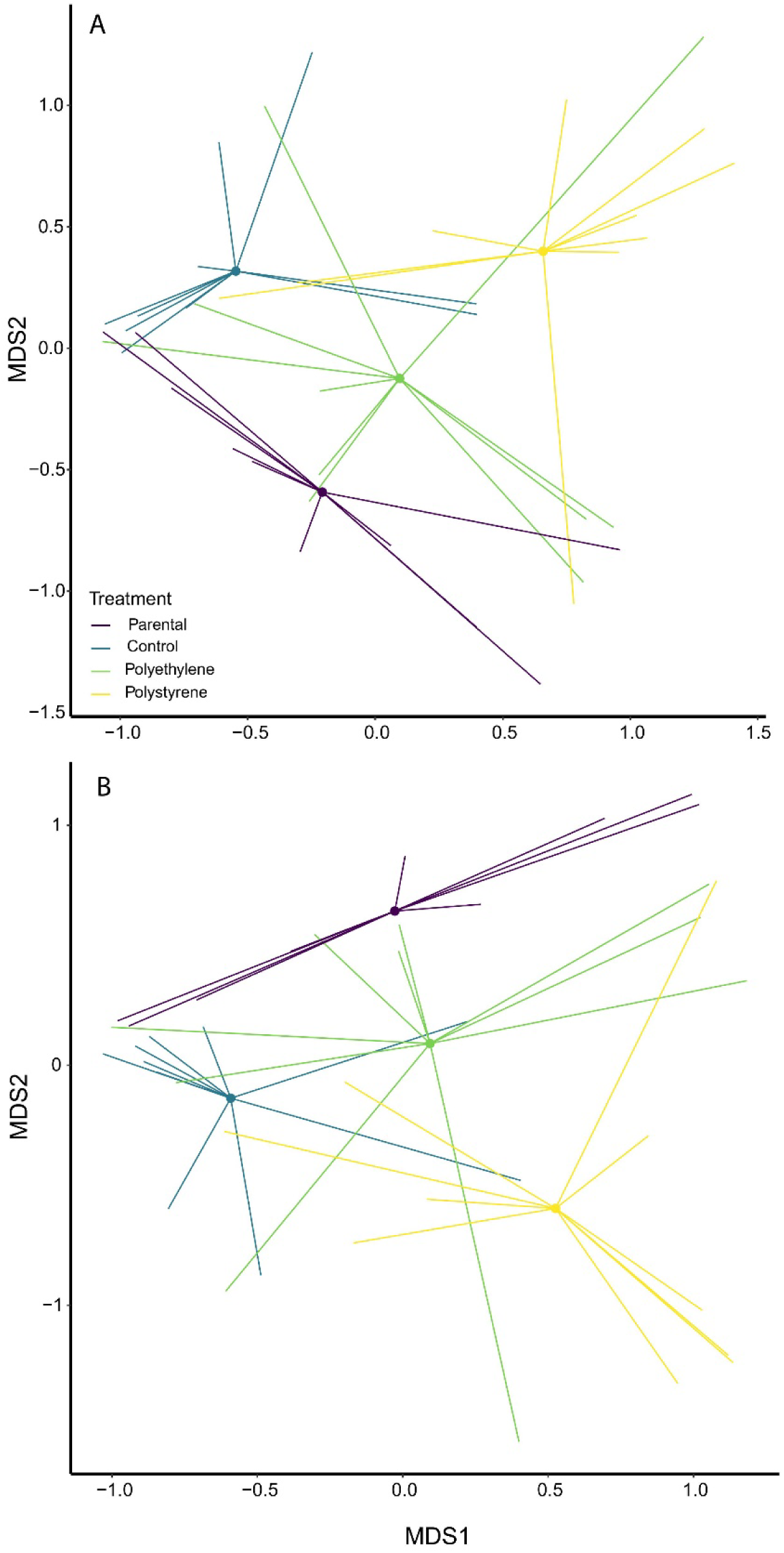
Non-multidimensional scaling (NMDS) plot based on Bray–Curtis distances of gastrointestinal microbiota in Tenebrio molitor larvae with (A) and without (B) Dickeya phage for the parental generation and the three treatments at the F_3_ generation. Lines represent individual samples originating from the centroid.

When the *Dickeya* phage was included, the Control and PS treatments from the F_3_ generation had similar Shannon-Weiner scores, which were higher than those of the parental generation, while the PE treatment was similar to the parental generation (Fig. 3A). Excluding the *Dickeya* phage, the median Shannon-Weiner score for PS decreased to a level similar to the parental generation, although the PS treatment retained high variance (Fig. 3B).

**Figure 3.**
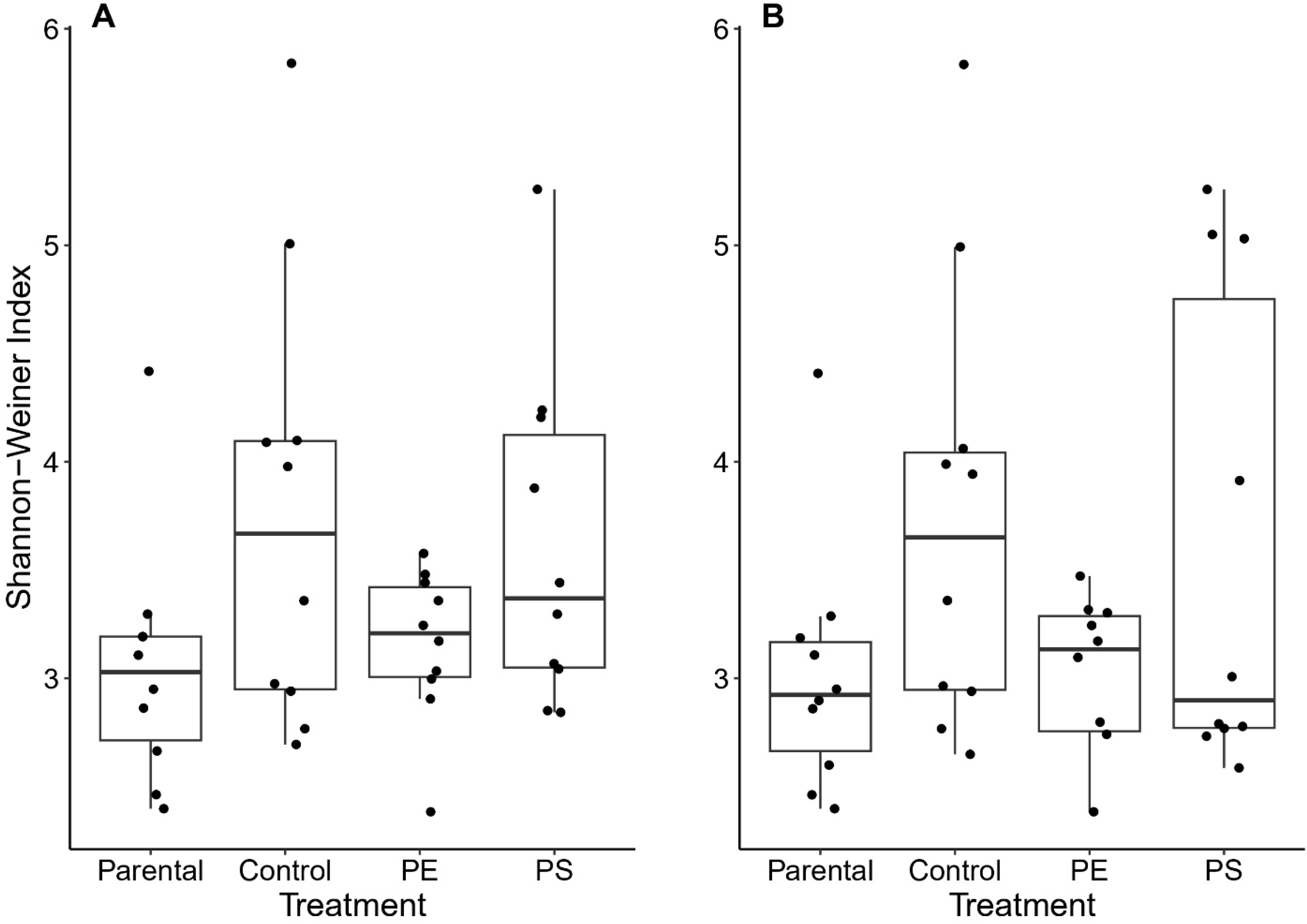
Shannon-Weiner indices with (A) and without (B) Dickeya phage for the parental generation and the three treatments at the F_3_ generation. Lines inside boxes indicate medians, boxes indicate 25th and 75th percentiles, bars indicate 1.5× the interquartile range, and points are individual measurements.

### Microbial Respiration

The incubations of gut microbes on PE and PS films yielded 30 strains. Fourteen were isolated from PE diets on PE cultures, five from control diets on PE or PS cultures and two from environmental controls on PE and PS cultures. Sequencing of 20 microbial isolates with the highest absorbance resulted in 17 unique species (Table 1; Fig. 4).

**Figure. 4.**
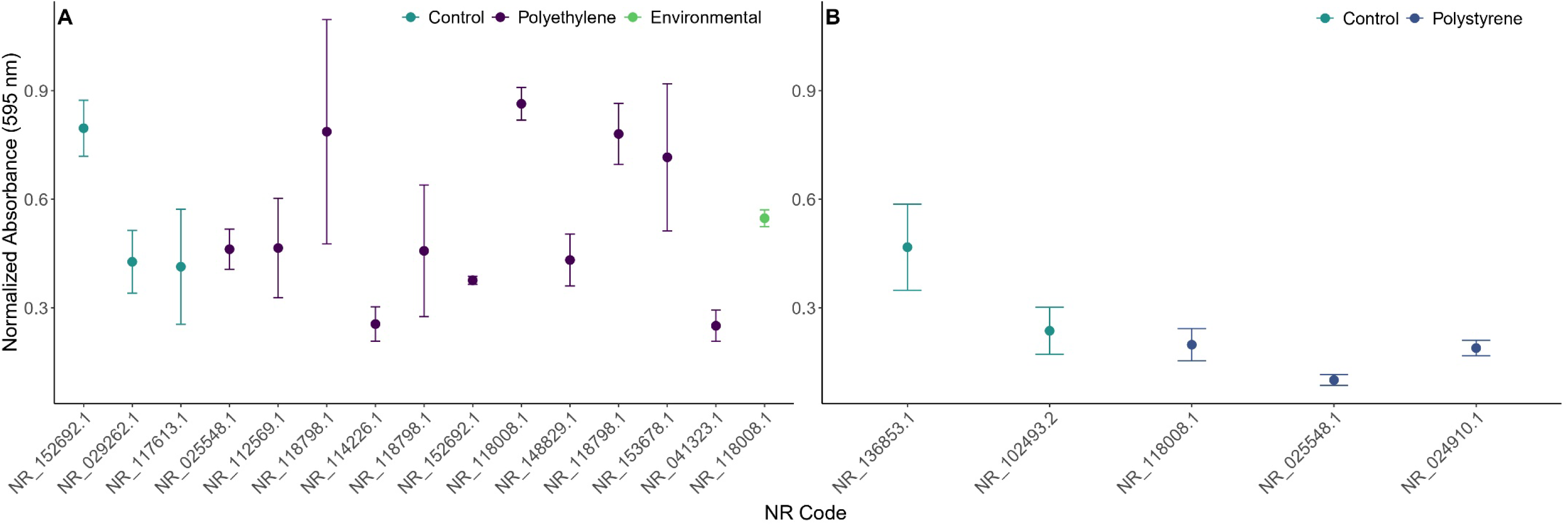
Microbial respiration of polyethylene (A) and polystyrene (B) microplastics by microbes isolated from Tenebrio molitor larvae fed wheat bran (control), polyethylene, and polystyrene.

**Table 1.**
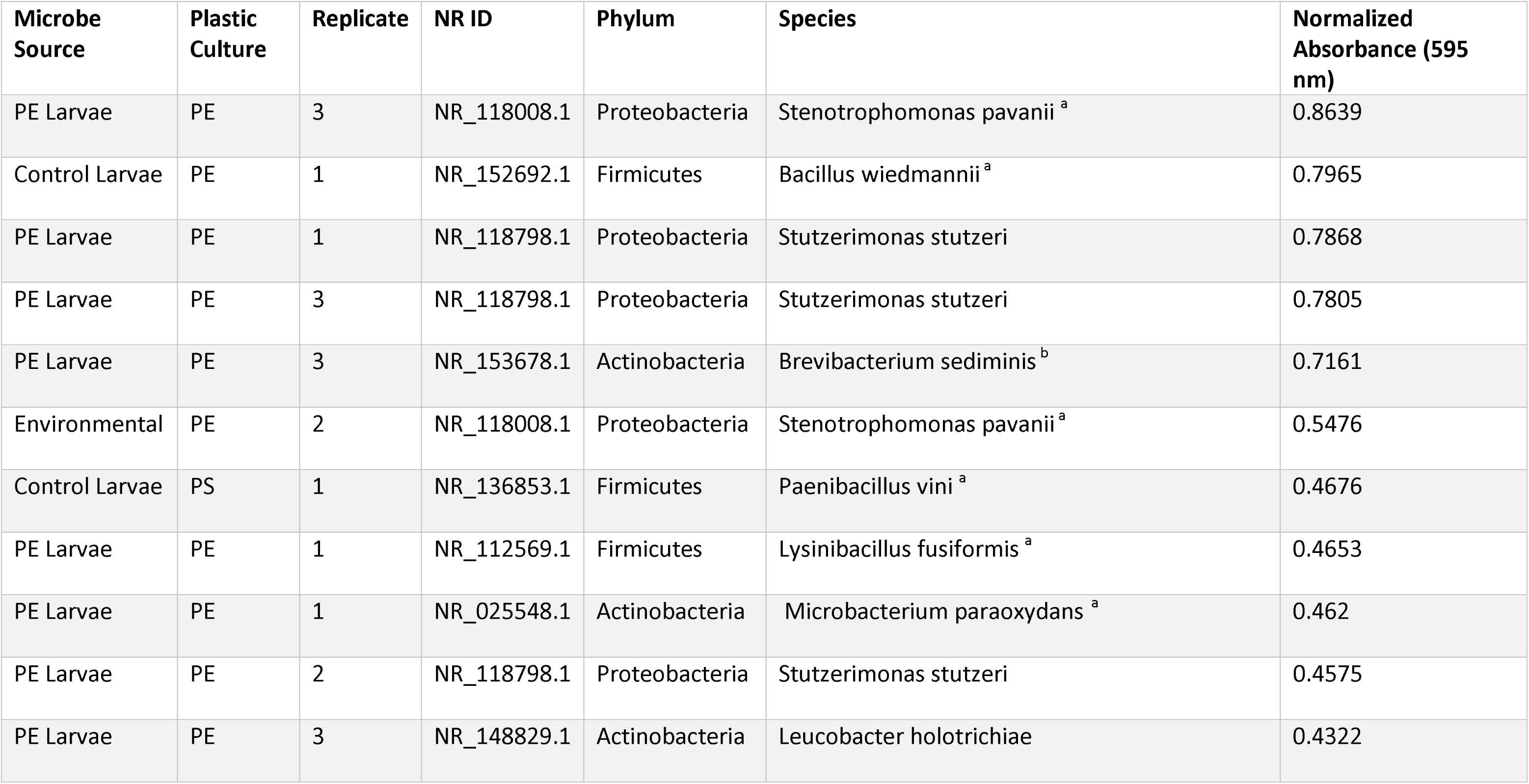
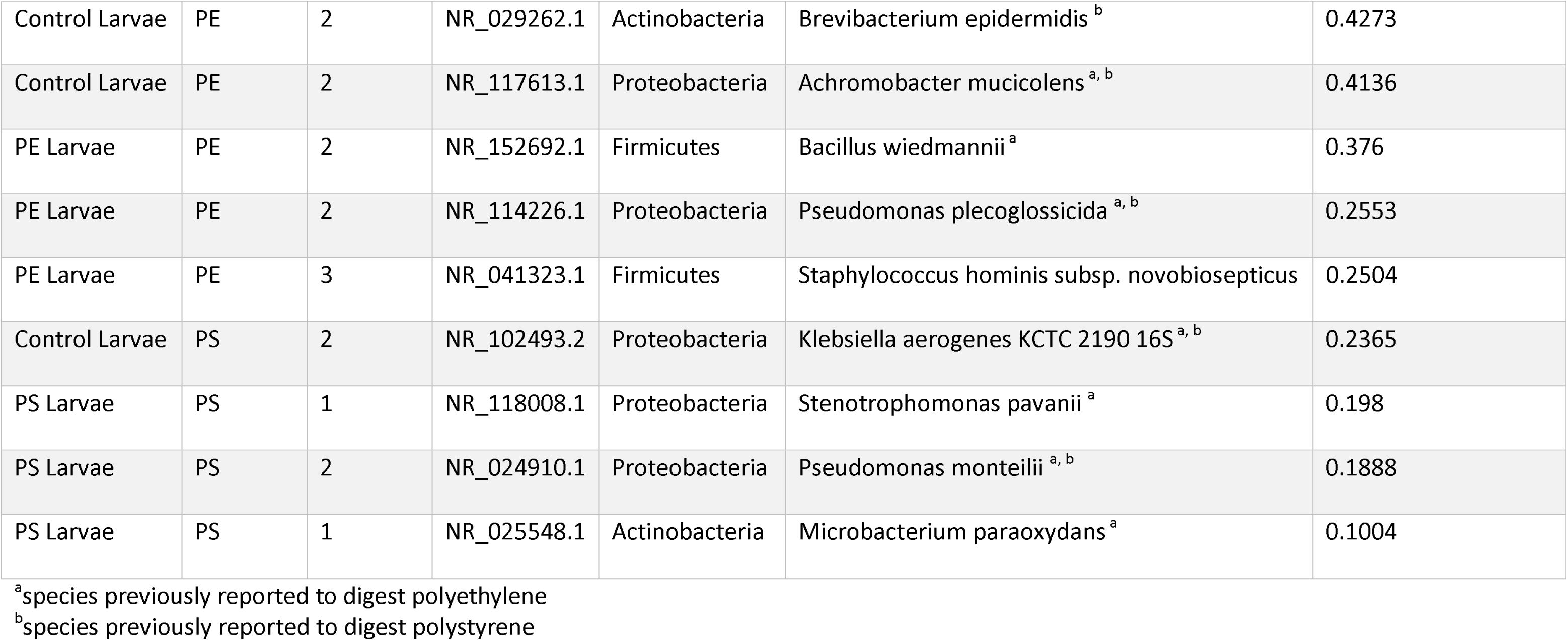
Sequenced plastic digesting bacterial isolates with corresponding respiration rate at 12 hours with the larval source of the microbes (microbe source), the plastic film it was cultured with (plastic culture), the flask replicate (Replicate), the NR identification (NR ID), the phylum (phylum), nearest species match (species), and the normalized absorbance measurement of plastic respiration (normalized absorbance).

| Microbe Source | Plastic Culture | Replicate | NR ID | Phylum | Species | Normalized Absorbance (595 nm) |
| --- | --- | --- | --- | --- | --- | --- |
| PE Larvae | PE | 3 | NR_118008.1 | Proteobacteria | Stenotrophomonas pavanii <sup>a</sup> | 0.8639 |
| Control Larvae | PE | 1 | NR_152692.1 | Firmicutes | Bacillus wiedmannii <sup>a</sup> | 0.7965 |
| PE Larvae | PE | 1 | NR_118798.1 | Proteobacteria | Stutzerimonas stutzeri | 0.7868 |
| PE Larvae | PE | 3 | NR_118798.1 | Proteobacteria | Stutzerimonas stutzeri | 0.7805 |
| PE Larvae | PE | 3 | NR_153678.1 | Actinobacteria | Brevibacterium sediminis <sup>b</sup> | 0.7161 |
| Environmental | PE | 2 | NR_118008.1 | Proteobacteria | Stenotrophomonas pavanii <sup>a</sup> | 0.5476 |
| Control Larvae | PS | 1 | NR_136853.1 | Firmicutes | Paenibacillus vini <sup>a</sup> | 0.4676 |
| PE Larvae | PE | 1 | NR_112569.1 | Firmicutes | Lysinibacillus fusiformis <sup>a</sup> | 0.4653 |
| PE Larvae | PE | 1 | NR_025548.1 | Actinobacteria | Microbacterium paraoxydans <sup>a</sup> | 0.462 |
| PE Larvae | PE | 2 | NR_118798.1 | Proteobacteria | Stutzerimonas stutzeri | 0.4575 |
| PE Larvae | PE | 3 | NR_148829.1 | Actinobacteria | Leucobacter holotrichiae | 0.4322 |
| Control Larvae | PE | 2 | NR_029262.1 | Actinobacteria | Brevibacterium epidermidis <sup>b</sup> | 0.4273 |
| Control Larvae | PE | 2 | NR_117613.1 | Proteobacteria | Achromobacter mucicolens <sup>a, b</sup> | 0.4136 |
| PE Larvae | PE | 2 | NR_152692.1 | Firmicutes | Bacillus wiedmannii <sup>a</sup> | 0.376 |
| PE Larvae | PE | 2 | NR_114226.1 | Proteobacteria | Pseudomonas plecoglossicida <sup>a, b</sup> | 0.2553 |
| PE Larvae | PE | 3 | NR_041323.1 | Firmicutes | Staphylococcus hominis subsp. novobiosepticus | 0.2504 |
| Control Larvae | PS | 2 | NR_102493.2 | Proteobacteria | Klebsiella aerogenes KCTC 2190 16S <sup>a, b</sup> | 0.2365 |
| PS Larvae | PS | 1 | NR_118008.1 | Proteobacteria | Stenotrophomonas pavanii <sup>a</sup> | 0.198 |
| PS Larvae | PS | 2 | NR_024910.1 | Proteobacteria | Pseudomonas monteillii <sup>a, b</sup> | 0.1888 |
| PS Larvae | PS | 1 | NR_025548.1 | Actinobacteria | Microbacterium paraoxydans <sup>a</sup> | 0.1004 |

<sup>a</sup>species previously reported to digest polyethylene
<sup>b</sup>species previously reported to digest polystyrene

As a group, microbes from larvae reared on a PE diet (0.50 ± 0.2 abs 595 nm) metabolized plastic more efficiently than those from control larvae cultured with PE (0.42 ± 0.2 abs 595 nm) at twelve hours. Strains isolated from the PS (0.16 ± 0.05 abs 595 nm) treatment had a lower level of respiration to those isolated from control larvae and cultured with PS (0.35 ± 0.2 abs 595 nm).

## DISCUSSION

After cultivating populations of *T. molitor* on a plastic and wheat bran diets for three generations, we found evidence that diets supplemented with PE and PS alter microbial community structure in *T*. *molitor*. We were also able to isolate and identify bacterial strains capable of plastic digestion, particularly those raised on and cultured with PE. Bacteria raised on and cultured with PS did not show increased plastic digestion above control.

*Spiroplasma* was found in high relative abundance in the Control (49%), PE (36%), and parental groups (43%), although nowhere near as high as previously reported 70-80% of relative abundance [38,41,52]. Studies have posited that *Spiroplasma*, is not only crucial for general digestion in larvae [53], but is a specialized strain maintained within *T. molitor* populations through lateral or hereditary means [38,52]. The genus was found in the lowest abundance in the PS treatment (<10%), which was unexpected as some suggest the *Spiroplasmataceae* family as important to PS degradation [54]. However, *Spiroplasma* may dominate early succession due to its initial colonization speed, rather than its efficiency at plastic digestion; Lou et al. [38] found that *Spiroplasma sp.* was dominant in both first (wheat bran only) and second succession (PS), but was replaced in the third succession (shift back to wheat bran only) by *Lactococcus*. We observed a similar shift where *Lactococcus* represented <1% in the parental generation, but was 12% of the control, and 6 and 7% of PE and PS respectively. *Lactococcus*is associated with the fermentation of carbohydrates and is an important genus in *T. molitor* digestion [38,54,55]. The shift between the wheat bran supplied by the commercial supplier and that of our lab could have selected for *Lactococcus*over multiple generations. Other genera also linked to general digestion in *T. molitor’s* included *Enterococcus* [36,38] and *Okibacterium* [37]. We found these genera in low abundances cumulatively representing less than 5% of the microbial community in the control and PS, while representing less than 1% in the PE and parental samples. This could be due to regional differences in the sourcing of larvae and/or wheat bran. Our phyla make up was similar to the results of other studies, both PS and PE treatments had high abundance of Firmicutes in their gut microbiome [37,38]. Proteobacteria was the second most abundant in PS samples, which also matched previous studies on *T. molitor* [37,54].

The composition and structure of the microbial communities change over time through stochastic and deterministic processes [56–58], The introduction of plastic into the diet has been shown to cause a shift in β-diversity when compared to a control diet [36,37], and when *T. molitor* transitioned off of plastic back to the original feed also had a lasting effect on the composition [38]. We saw shifts in β-diversity for both PS and Control diets, while the PE continued to resemble the parental microbial make up which contrasts results from previous studies [36–38,59]. This result can be interpreted several ways: first, the addition of PE was not impactful enough to cause a deterministic shift; second, the PE treatment was able to maintain the existing microbial assembly and specialize existing strains; third, the Control larvae experienced a stochastic shift that resembled the PE treatment.

The addition of PS in the diet may have induced dysbiosis, a disruption in the diversity, abundance or function of the microbiota that affects host physiology. This potential dysbiosis would have left the larvae in that treatment susceptible to colonization by *Dickeya* phage. The Shannon-Weiner scores with *Dickeya* phage removed indicate lower microbial diversity in the PS treatment, relative to the PE treatment. PS is also a more complex polymer and more difficult to digest [4,18]. In the PS treatment, the *Dickeya* phage represented 30% of the relative abundance. *Dickeya* phage is a bacteriophage of *Dickeya spp.*, which is a known plant pathogen, specifically linked to soft rot and blackleg of many potato varieties [60]. Additionally, *Dickeya* spp. have been found to cause septicemia in some insect species [59,61]. In normal circumstances the presence of a phage would accompany the presence of its associated bacteria [62], however *Dickeya spp.* were absent from our 16S data. Phage-gut bacteria interactions in insects is not a well-explored topic, however T-phages, a group of bacteriophages that infect *Escherichia coli*, may play a key role in shaping a host’s microbial community [63,64]. Additionally, indirect positive effects on other bacterial abundances have been seen in some insects infected with a genus specific phage [65]. The entry vector of the *Dickeya* phage was likely the potatoes given to all treatments as their source of moisture. The higher relative abundance of *Dickeya* phage in PS larvae suggests that the complexity of PS may have hindered its digestion, disrupted the gut microbiome of *T. molitor*, and created an environment favorable to phage proliferation. However, further exploration is needed to clarify the specific interactions between *Dickeya* phage and the host microbiome.

Due to the inherent pseudoreplication in our experimental design—where all larvae within each treatment were co-housed—the observed differences could be attributed to either the treatment effects or the co-housing of larvae. Since the samples are not truly independent, the p-values obtained from the PerMANOVA may not accurately reflect the true variability between treatments. Therefore, our results should be interpreted with caution. However, we think it is likely that the effect of diet is responsible for our observed differences between treatments, as the effect of co-housing was probably weak in comparison, as diet is a strong driver of microbiota community structure [66–69]. The treatments were housed side-by-side, originated from the same source, handled by the same people, and fed the same wheat bran and potatoes. Any bacterial species would have had sufficient opportunities to colonize all three treatments and the interactions between individual larvae do not seem likely to cause the observed differences.

We isolated 30 morphotypes of plastic digesting bacteria across all treatments. Of the five highest metabolic rates, four strains came from the PE cultures, with the last from a control on PE culture. We were able to broadly exhibit that generational improvement of plastic digestion in PE isolates compared to control. PS samples did not see the same generation improvement, as isolates originating from control larvae had a higher average metabolism compared to those from PS larvae. We found 17 unique plastic digesting strains after sequencing our top 20 samples, nine of which were previously linked to digesting PE plastics, while six were associated with PS. A previous association between PS and the *Brevibacterium* genus has been established, however we isolated two unique strains: one from a PE and one from a control culture [36,70]. We additionally identified three strains that were not previously associated with plastic digestion; *Stutzerimonas stutzeri*strain CCUG 11256 (associated with heavy metal phytoremediation) [71], *Leucobacter holotrichiae* strain T14 (previously identified in *Holotrichia oblita*) [72], and *Staphylococcus hominis subsp. novobiosepticus* strain GTC 1228 (anti-biotic resistant strain first sampled in human blood) [73], all of which were isolated from PE samples. *Stutzerimonas stutzeri*was previously classified in the *Pseudomonas* genus, which includes several species which have been found to digest plastic [54], but not the specific strain we found.

While evolution *in situ* may have occurred and promoted better plastic digestion in general, a strain-to-strain comparison suggests that may not be the most important factor. *Bacillus wiedmannii* strain FSL W8-0169 of the *cereus* group was isolated and identified in both a control and PE sample, where the control sample had a higher respiratory rate (0.78) when compared to the PE sample (0.37). There was also a strain of *Brevibacterium* sp., a genus previously shown to digest PE [36,70], found again in both control (0.40) and PE (0.94). All three of the PE cultures produced a plastic digesting *S. stutzeri*, and the respiration rates varied from 0.40-0.79. One of the PE culture replicates produced five unique strains; three of which were among the five strains with the highest respiration rate, indicating that plastic digestion is not only strain dependent but also sample dependent. This may also imply a synergistic effect between strains. Further evidence of synergistic effects is apparent when comparing the identified strains from PEPE3, the *Brevebacterium sp.*isolated from this sample was more efficient than the other *Brevebacterium sp.* This was also the case for the *Stenotrophomonas pavanii* strain LMG 25348, which was also detected in a PS sample as well as one of three PE process controls. This strain was most efficient from the PE culture with a respiration rate of 0.86, where the PS and process control saw rates of 0.19 and 0.54 respectively. The presence of *S. pavanii* in a process control likely arose through aerosol contamination from a PE sample during a broth change as it was only isolated from three of the eighteen gut cultures. If the contamination was from an air or environmental source, we would expect to have isolated this strain from all cultures and see a more uniform distribution of plastic digesting rates. *Stenotrophomonas pavanii* was initially characterized in 2011 as a nitrogen fixing bacteria in sugar cane [74]. The example of *S. pavanii* further suggests that the PS larvae were unable to develop specialized strains for breaking down PS despite multiple generations exposed to the plastic. There is a possibility that the PS microbes under preformed due to the transition from selective media (BH broth) to non-selective media (TSA plates). Follow-up work should account for this by comparing metabolism on BH broth relative to TSA. The intra-strain metabolic differences we observed highlight the need for more technical replicates in subsequent studies to parse if plate-specific differences arise from varying starting bacterial communities or *in vitro* evolution. Additionally, to explicitly test for synergistic interactions, future work should examine the plastic metabolism of gut-derived bacterial isolates in a community and in isolation.

In conclusion, we found that long-term polyethylene (PE) supplementation led to an improvement in plastic digestion by gut bacteria isolated from *T. molitor* larvae, whereas polystyrene (PS) supplementation did not yield the same results. Specifically, multiple bacterial species capable of metabolizing PE were identified, suggesting a possible synergistic effect within the gut microbiome that enhanced plastic degradation. This improvement, however, was strain– and sample-dependent as not all microbial isolates had the same capacity for plastic digestion The lack of a clear improvement in PS degradation and potential signs of compromised gut health in PS-fed larvae may indicate limitations of our approach for more recalcitrant plastics. However, the presence of *Dickeya* phage in the PS treatment group suggests possible dysbiosis in the gut microbiota, which may have hindered microbial plastic metabolism. Further research could clarify the mechanisms behind microbial plastic digestion and to explore whether *in situ* selection or *in vitro* culturing exerts a greater influence on improving degradation efficiency. Additionally, while PE showed promise for enhanced degradation, more work is needed to determine whether these microbes can be eventually useful for practical applications. Future research could measure the amount of degradation of micro– and nanoplastics in culture and use Fourier transform infrared spectroscopy to assess degradation of macroplastics.

## Author Contributions

**Hannah McKinnon Reish**: Formal analysis, investigation, methodology, validation, visualization, writing-original draft preparation, writing – review and editing; **Rebecca F. Witty**: Investigation; **Adam H. Quade**: Formal analysis, visualization; **Jason W. Dallas**: Formal analysis; writing – review and editing; **Lucas J. Kirschman**: Conceptualization, data curation; formal analysis, funding acquisition, methodology, project administration, resources, supervision, validation, visualization, writing – review and editing.

