## Supplemental Tables 1 & 2 for "Isolation of Plastic Digesting Microbes from the Gastrointestinal Tract of *Tenebrio Molitor*"

| Table S1. Results of the PERMANOVA using rarified gut microbiota community data that includes the bacteriophage with the Bray-Curtis dissimilarity matrix and 10,000 permutations. | | | | | |
| --- | --- | --- | --- | --- | --- |
|  | **Df** | **SOS** | **R^2^** | **F** | **Pr(>F)** |
| **Dietary Treatment** | 3 | 2.80 | 0.22 | 3.29 | 4e-04 |
| **Residual** | 36 | 10.23 | 0.78 | -- | -- |
| **Total** | 39 | 13.03 | 1.00000 | -- | -- |
| Table S2. Results of the PERMANOVA using rarified gut microbiota community data with the bacteriophage removed with the Bray-Curtis dissimilarity matrix and 10,000 permutations. | | | | | |
|  | **Df** | **SOS** | **R^2^** | **F** | **Pr(>F)** |
| **Dietary Treatment** | 3 | 2.61 | 0.20 | 2.99 | 0.0007 |
| **Residual** | 36 | 10.49 | 0.80 | -- | -- |
| **Total** | 39 | 13.11 | 1.00000 | -- | -- |
